# Refining a molecular tool kit to capture tropicalization in Mediterranean Marine Protected Areas

**DOI:** 10.1101/2025.05.26.656120

**Authors:** Erika F. Neave, Charalampos Dimitriadis, Peter Shum, Ana Riesgo, Stefano Mariani

## Abstract

Tropicalization, the process by which tropical species expand their ranges poleward due to global ocean warming, is a prominent threat to Mediterranean marine ecosystems, challenging their effective management and conservation. The arrival of non-indigenous fish is exacerbated by the Suez Canal that allows Indo-Pacific species to enter the region from the Red Sea. While some non-indigenous species (NIS) have already changed the composition and function of native Mediterranean communities, many others continue to arrive but often go unnoticed without regular surveys or until they are strongly established. We conducted biodiversity surveys using underwater visual census (UVC), aqueous environmental DNA (eDNA) and sponge-derived eDNA (i.e., eDNA accumulated in sea sponge tissue) at two protected locations in Zakynthos, Greece. Seven NIS were detected by eDNA, but only one (dusky spinefoot, *Siganus luridus*) was detected with all three methods. The fish assemblages of the two locations could be distinguished based on both UVC and eDNA but not from sponge-derived eDNA data, perhaps given low filtration rates. Of the three methods, aqueous eDNA metabarcoding provided the most comprehensive species list including new NIS detected (with the red-toothed triggerfish, *Odonus niger*, and the houndfish, *Tylosurus crocodilus*, being the first and third record in the Mediterranean Sea, respectively). Our findings highlight the potential value of incorporating molecular methods into regular monitoring as early warning tools for detecting NIS in marine protected areas threatened by ocean warming.

## Introduction

With the current pace of climate change temperate habitats are prone to tropicalization, a phenomenon defined by poleward distribution shifts of warm-affinity species in response to sea warming (McLean et al., 2021; Osland et al., 2021; Vergés et al., 2014). The Mediterranean Sea is undergoing accelerated warming compared to other ocean basins (Urdiales-Flores et al., 2023), resulting in an influx of non-indigenous species (NIS) (Lejeusne et al., 2010). In addition to warming, tropicalization is amplified in this region due to the Suez Canal (1869), which provides an artificial waterway for NIS to travel from the Red Sea to the Mediterranean (Katsanevakis et al., 2014). Over 500 tropical NIS have entered via the canal and these species are concentrated in the eastern Mediterranean (Galanidi et al., 2023; Edelist et al., 2013; Katsanevakis et al., 2014; Raitsos et al., 2010). Warming sea temperatures, facilitating NIS in this region, have caused substantial shifts in the structure of native communities and the deterioration of habitats and ecosystem functioning (Peleg et al., 2020; Vergés et al., 2014; Zarzyczny et al., 2023), resulting in mixed effects on ecosystem services such as biological resources (e.g., fisheries, aquaculture) and human health (Tsirintanis et al., 2022). At present, the Mediterranean harbours around 1000 NIS and newly established species increased by 40% between 2011 and 2021 (Zenetos et al., 2022). Tropicalization is causing rapid changes, such that monitoring of NIS is frequently acknowledged as a crucial priority for effective environmental and socioeconomic management of the Mediterranean (i.e., EU Regulation 1143/2014 for alien species; European Commission. Joint Research Centre., 2021).

Generating cost-effective, reliable biodiversity evidence remains a challenge for environmental managers, especially at sea. Underwater visual census (UVC) is a standard method for collecting biodiversity information in Mediterranean Marine Protected Areas (MPAs) (Giakoumi et al., 2017). However, this approach is time consuming, requires the expertise of skilled taxonomists, and introduces observer bias, which is difficult to mitigate over the timescales relevant to long-term monitoring. Environmental DNA (eDNA) is complementary to UVC, with higher sensitivity to detect more taxa, such as cryptobenthic (Bessey et al., 2023), highly elusive pelagic species (Bakker et al., 2017) and NIS (Jerde et al., 2011). While eDNA analyses are becoming more cost-effective, they have not been routinely incorporated into coastal monitoring strategies. This is often due to unfamiliarity with the technology or uncertainty on whether the data can complement traditional visual information at managed sites. Visual census offers more explicit quantitative data, such as abundance information, which managers are generally more accustomed to relying on. However, several countries are beginning to adopt eDNA analyses in monitoring strategies and biosecurity measures (De Brauwer et al., 2023; Jerde et al., 2013; Stepien et al., 2022.).

Mediterranean MPAs are vulnerable to biological invasions, even after only a few years from the establishment of NIS (Dimitriadis et al., 2024), with tropicalization acting as an exogenous threat that challenges conservation efforts and outcomes (Giakoumi et al., 2017). Tropicalization causes functional shifts in fish communities leading to ecosystem phase shifts that can have effects on behaviours and genotypic composition of species (Vergés et al., 2014; Zarzyczny et al., 2023). These problems have been identified as key conservation issues in Zakynthos MPA (Giakoumi et al., 2019) and other Greek islands with similar habitats (Bianchi et al., 2014). Here we conducted biodiversity surveys at two MPAs comparing traditional UVC methods to aqueous eDNA. We also collected biopsies from marine sponges, filter-feeding organisms that have recently been found to collect eDNA, an approach termed ‘natural sampler DNA’ (nsDNA) (Mariani et al., 2019; Turon et al., 2020). Metabarcoding of marine sponge nsDNA has been shown to be highly comparable to aqueous eDNA for detecting fish species of various sizes in artificial (Cai et al., 2023) and natural environments (Jeunen et al., 2021). Together with conventional UVC methods we explored the potential for molecular sampling strategies to expand the tool kit that managers can use to monitor marine sites of ecological and cultural significance.

## Methods

### Study Area

Two sampling locations within the marine protected areas of Zakynthos (Eastern Ionian Sea, Greece), were monitored for fish biodiversity using UVC, eDNA, and sponge nsDNA (Figure 1). Korakonissi (Figure 1C), within the protected area of 92/43/EU Directive under code GR2210001, is a popular swimming location consisting of rocky reefs, gravel and sand substratum with depths between ∼4-12 m where we sampled. This area has low protection status, with no specific measures for fish conservation in effect. While Dafni beach (Figure 1B), has the maximum protection level (i.e., no-take, no-go from May to October, only swimming is allowed) within the National Marine Park of Zakynthos, and consists of a mixture of shallow rocky reefs, seagrass beds and sand substrate habitats. At each location, visual census surveys (Supplementary Text), aqueous eDNA and sponge collection occurred between September 10-14, 2021 (Figure 1A).

**Figure 1.**
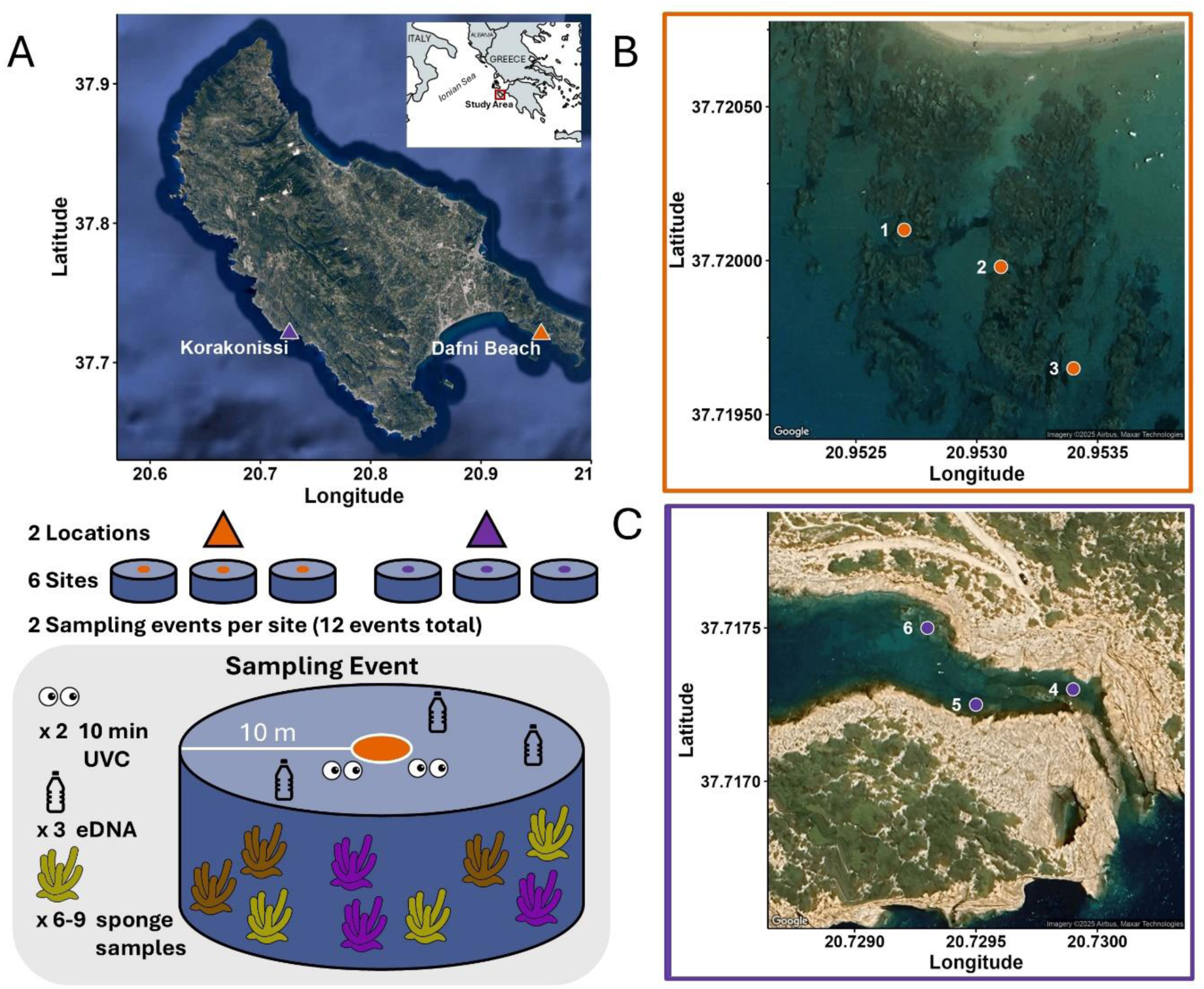
A) Map of Zakynthos, Greece showing the two sampling locations (orange and purple triangles) and a diagram of the sampling design. The six sampling sites (orange and purple points) were sampled twice each from the locations: **B)** Dafni Beach, National Marine Park of Zakynthos and **C**) Korakonissi.

### Aqueous eDNA Sampling and Sponge Collection

Two snorkelers conducted underwater visual census (UVC) (Bohnsack & Bannerot, 1986), while seawater was collected by other snorkelers in 2 L bottles for eDNA metabarcoding analysis. Sample bottles were filled 1 meter below the surface. Filtration blanks (2 L of filtered bottled drinking water each) were performed onshore where samples were filtered and stored, before and at the end of each sampling day (n = 8), while field blanks (also 2L each) were collected at each location twice, once per sampling day (n = 4) (Supplementary Text). Each triplicate sample was obtained from different areas within a 10 m radius of the sampling point, to get an average representation of each sampling point (two locations X three sites X three replicates X two sampling events: n = 36) (Figure 1A). The seawater was pushed through a 0.45 µm Sterivex filter (PES membrane, Merck Millipore) using a 60 mL syringe (Fischer Scientific). Filters were then sealed into individual bags. Where possible triplicate biopsies (∼2 cm^3^) of two to three sponge species per sampling point (four unique species total) were taken, cutting into the inner cortex of the sponge using a dive knife and pliers, at each site and sampling event (n = 85) (Table 1). Sponge biopsies were stored in 15 ml falcon tubes containing 100% molecular grade ethanol. Filters and sponges were stored in a cool box before transfer to −20°C at the end of each sampling day.

**Table 1.**
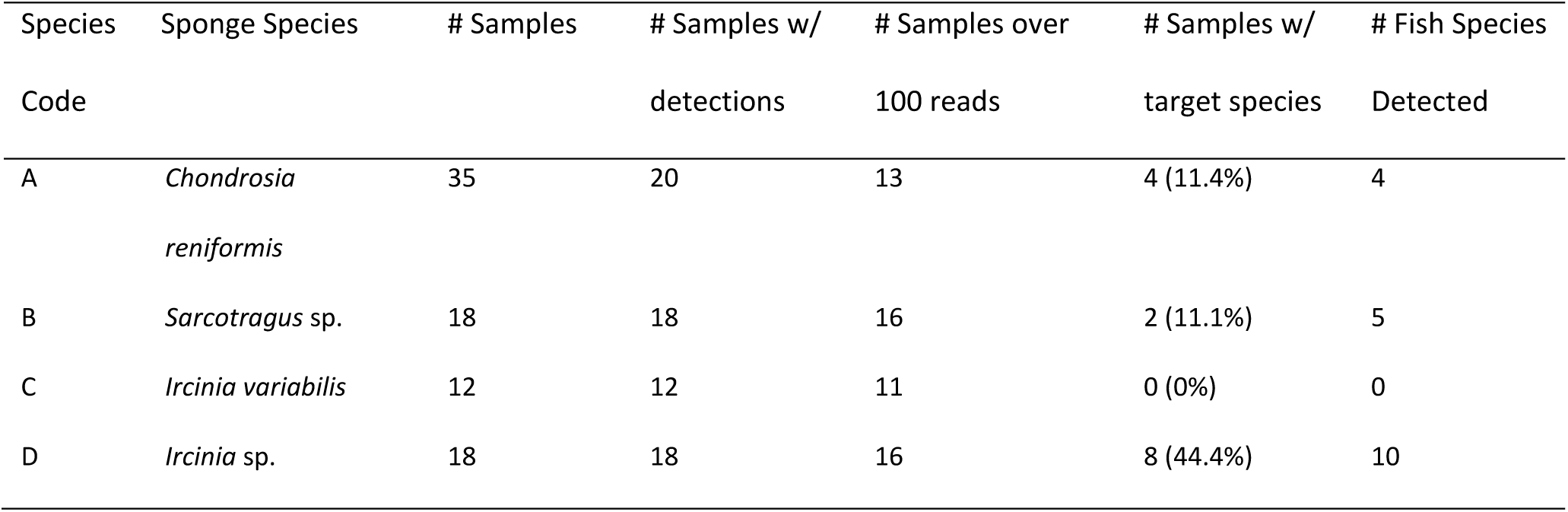
Sponge species tested as candidates for sponge-derived eDNA.

### DNA Extractions

The DNA extractions were performed in a designated eDNA lab, separated by two doors from any other lab, and where users must wear hazmat suits, masks and caps to mitigate contamination. Equipment was washed in a 10% bleach solution, followed by 5% Lipsol detergent then Milli-Q water, and exposed to ultra-violet light for 30 minutes. DNA was extracted from the sterivex filters using a modular universal DNA extraction protocol ([version 2](https://www.protocols.io/view/mu-dna-a-modular-universal-dna-extraction-method-a-6qpvryj2gmkn/v2)); specifically, all steps of the Mu-DNA tissue extraction protocol with the addition of an inhibitor removal step taken from the mu-DNA soil protocol, which resulted in 100 μl of DNA extract (Sellers et al. 2018). Sponge tissue was dried by blotting excess storage ethanol onto filter paper inside a petri dish using tweezers (Harper et al., 2023). Between 0.25 and 0.35 grams dry-weight of sponge were placed with lysis buffer into a 1.5 ml microtube and incubated at 55°C for 24 hrs. DNA was extracted from the sponge tissue using the same modified Mu-DNA tissue protocol used for the sterivex filters (Harper et al., 2023; Sellers et al., 2018). An extraction blank was processed with each batch of filter and sponge DNA extractions (n = 3, n = 4, respectively). Sponge identification was aided by Sanger sequencing (Supplementary Text).

### eDNA Library Preparation and Sequencing

Sponge DNA extracts were diluted with molecular grade water to between 30-50 ng/µl, while aqueous eDNA extracts were unaltered (all < 5 ng/µl). DNA was PCR amplified using the Tele02 8-bp dual barcoded primers (Taberlet et al., 2018): Tele02-F (5’-AAACTCGTGCCAGCCACC-3’) and Tele02-R (3’-GGGTATCTAATCCCAGTTTG-5’), which were used to target a ∼167 bp fragment of the mitochondrial 12S ribosomal RNA gene. PCRs were prepared to a total volume of 20 μl for each sample and included 10 μl of 2X MyFi Mix (Meridian Bioscience), 1 μl of each forward and reverse primer (10μM), 0.16 μl Bovine Serum Albumin (Thermo Fisher Scientific), 5.84 μl molecular grade water, and 2 μl of DNA extract. All samples were amplified in triplicate using the following conditions: 95°C for 10 min, followed by 35 cycles of 95°C for 30 s, 60°C for 45 s, 72°C for 30 s, and a final extension at 72°C for 5 min followed by a 4°C hold. Negative controls (molecular grade water, n = 6) and positive controls (extract of iridescent catfish, *Pangasionodon hypopthalmus*, which is not present in the Mediterranean, n = 6), underwent PCR alongside the samples. PCR triplicates were visualized on a 2% agarose gel stained with SYBRsafe dye and pooled. PCR products were individually purified twice using a 1:1 followed by 0.6:1 ratios of Mag-Bind® Total Pure NGS magnetic beads (Omega Bio-Tek) to PCR product. Products were visualised on an agarose gel again to assure purity (i.e., target length bands on agarose gels were visible with minimal to no other bands present). Purified PCR products were quantified using a Qubit dsDNA HS Assay kit (Invitrogen) and pooled at equimolar concentration into separate aqueous eDNA and sponge nsDNA libraries. The size integrities of the libraries were analyzed on a TapeStation 4200 (Agilent). The libraries were then purified based on the TapeStation results by repeating the magnetic beads procedure with the same ratios explained before. A unique adapter sequence was ligated to each library using the NEXTFLEX® Rapid DNA-Seq Kit for Illumina (Revvity) following the manufacturer protocol. After adapter ligation, the libraries visualized on the TapeStation and purified with magnetic beads with a 0.8:1 ratio of beads to sample. The dual-indexed libraries were then quantified by qPCR using the NEBNext® Library Quant Kit for Illumina (New England Biolabs) following the manufacturer protocol prior to being sequenced on an Illumina iSeq100 (Supplementary Text).

### Statistical Analysis

Prior to the analysis, the sequences were processed using a standard bioinformatics pipeline (Supplementary Text). Statistical analyses were done using R version 4.1.3 (R core team 2022). Any contamination present in controls was removed 10-fold from the samples to decrease the likelihood of false positives (i.e., 1 read of *Thalassoma pavo* was present in a negative control, so 10 reads of *Thalassoma pavo* were removed from each sample in the dataset) (Supplementary Table 1). This strict decontamination protocol was followed since we were particularly interested in non-indigenous species detections but wanted to be conservative in our conclusions. For management purposes, false positives could be more favorable than false negatives, and this should be considered when designing a decontamination protocol. For downstream analysis, only genus or species level assignments were retained. Any sample with less than 100 reads was removed from the dataset based on the assumption that this threshold indicated the sequencing was unsuccessful, since the samples removed had a factor of ∼10-100 less reads than the remaining samples. This threshold removed 71 (84% of total) sponge samples and 25 (69% of total) aqueous eDNA samples, of which 53 sponge samples and 16 aqueous eDNA samples had zero reads. While we believe the sample loss for the aqueous eDNA samples was due to an aggressive inhibitor removal step in the extraction process that is typically used for soil samples, the lack of success for certain sponges is likely due to PCR inhibition (Cai et al. 2022). Loss of some aqueous eDNA samples were further analyzed by comparing this study to the only other study to our knowledge which includes environmental DNA samples from Zakynthos (Aglieri et al., 2021). Aglieri and colleagues’ data was cleaned to retain species level detections with a percent identity ≥ 98%, the same threshold used in our study, so that the datasets could be directly compared. Species richness, Shannon Index and J-evenness for each method (i.e., eDNA, sponge, UVC) were calculated using the BiodiversityR package v 2.14.2.1 (Kindt, 2023). The J-evenness was calculated by dividing the Shannon Index by the log of species richness (Pielou, 1966). These diversity indices for each method were compared at each location using Mann-Whitney U (Wilcoxon rank-sum) tests or Kruskal-Wallis with a post-hoc pairwise Wilcoxon rank-sum test.

The R package vegan v 2.5.7 (Oksanen et al., 2009) was used to compare beta-diversity between sampling locations. Jaccard dissimilarity indices were calculated from genus and species-level presence-absence data, resulting in separate matrices for UVC and aqueous eDNA observations. Matrices were also made of Bray-Curtis dissimilarity, calculated from Hellinger transformed count and read data (i.e. UVC and aqueous eDNA, respectively). Non-metric multi-dimensional scaling (NMDS) plots were created for each distance matrix. Stress plots of each ordination were observed and led to the removal of three outlier eDNA samples from Korakonissi (Supplementary Figure 1, 2). The following tests were performed on both Jaccard and Bray-Curtis indices (Supplementary Figure 3): homogeneity among the group dispersions of locations (i.e. Dafni Beach and Korakonissi) and for each sample type, differences in beta-diversity between locations were tested with a permutational multivariate analysis of variance (PERMANOVA). The sampling location map was made with the following R packages: rstudioapi v 0.15.0 (Ushey et al., 2024) and ggmap v 3.0.2 (Kahle & Wickham, 2013). All remaining figures were generated using the R packages tidyverse v 1.3.1 (Wickham et al., 2019) and ggplot2 v 3.4.0 (Wickham, 2011), except for Figure 3 which was created in Adobe Illustrator.

## Results

### Aqueous eDNA detections and UVC observations

Both eDNA and sponge sequencing runs were combined for a total of 536 molecular operational taxonomic units (MOTUs) assigned to the bony fishes (class: Actinopterygii). Of those, 417 MOTUs assigned to genus and species level taxonomy. The taxonomic assignments were filtered to a likelihood of identity, in this case ≥ 98% identity, and resulted in matching assignments that could be merged into 27 species and 14 genera (41 taxa total from eDNA and nsDNA combined) (Supplementary Table 2). When combining the UVC data with eDNA data, there were 55 taxa detected or identified in total (Figure 2) of which 39 were at the species-level (Figure 3). When combining molecular data (41 taxa) the sponge nsDNA did not contain unique detections (Figure 4), so indeed 55 taxa were identified when considering all three methods. At Dafni Beach 24 taxa were detected by eDNA and UVC, seven of which were detected by both methods. At Korakonissi, 33 taxa were detected by eDNA and 22 by UVC, with 10 being detected by both methods (Figure 2).

**Figure 2.**
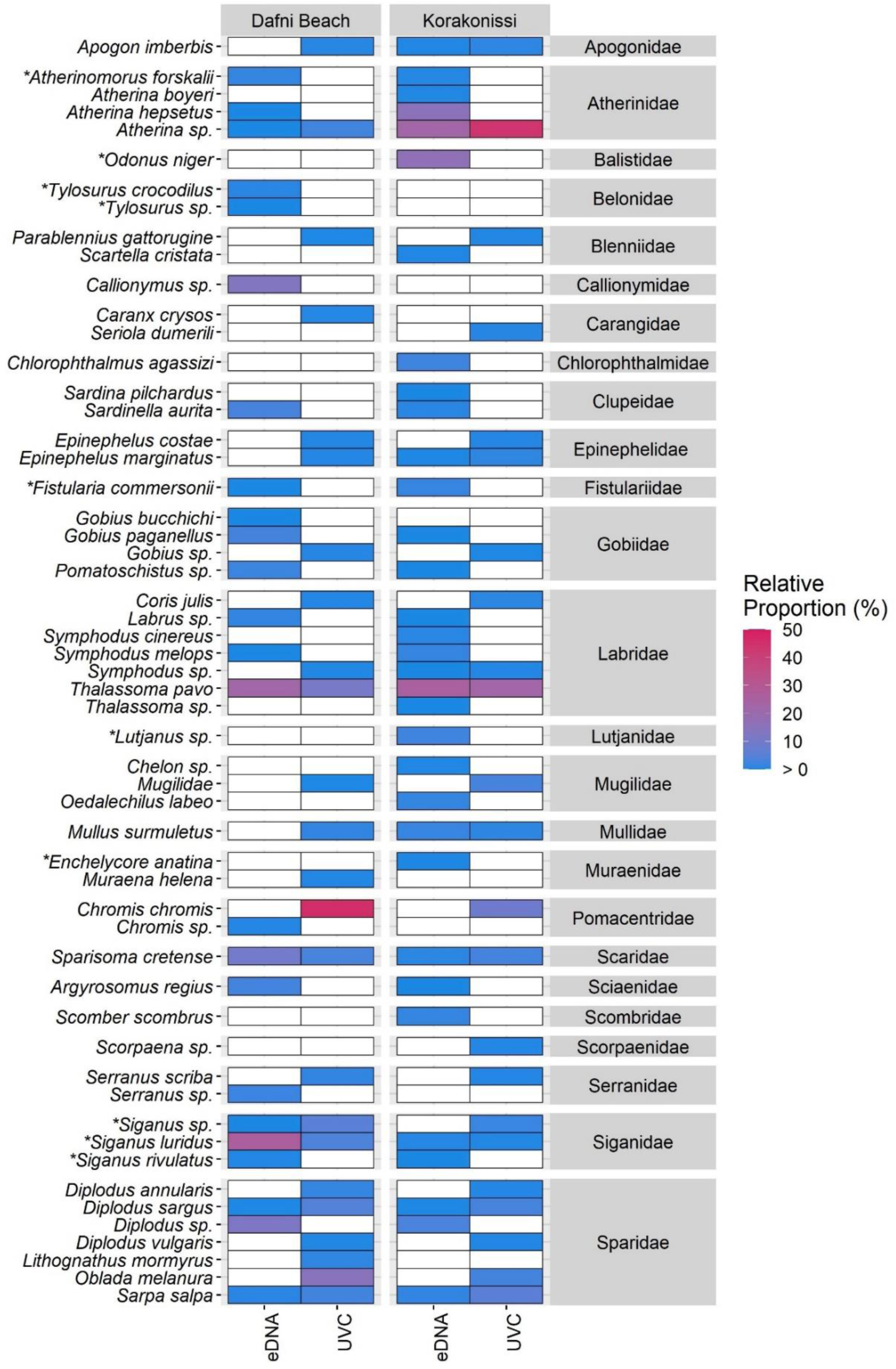
Heatmap showing the relative proportion of detections (eDNA) and counts (UVC). (*) Non-indigenous species.

In some instances, the eDNA data helped improve species-level detection. One example of this was evidence of non-indigenous rabbitfish species (genus: *Siganus*). Despite the known presence of two species of *Siganus* in the area, observers could only confidently recognize *Siganus luridus*, recording the remaining rabbitfish observations as *Siganus* sp. at both locations. At Dafni Beach the largest proportion of eDNA detections was of *Siganus luridus* (26.4%; 13,731 reads), while 0.16% (82 reads) belonged to *Siganus rivulatus*. Both *Siganus* species were also detected at Korakonissi making up 0.46% of eDNA read counts. Another instance where eDNA provided greater taxonomic granularity than UVC was for the small schooling sand smelts (Family: Atherinidae). At Korakonissi, *Atherina* sp. made up 44.1% of the visual counts, while eDNA from the same location detected three species from the family Atherinidae (i.e., *Atherina boyeri*, *Atherina hepsetus* and *Atherinomorus forskalii*), where their combined relative proportion including the *Atherina* sp. genus-level assignment was 38.4%.

However, there were also cases where UVC had greater taxonomic resolution than eDNA metabarcoding. Three species of sparidae in the genus *Diplodus*, were identified by UVC (i.e., *Diplodus annularis*, *D. sargus* and *D. vulgaris*) while only one species, *Diplodus sargus*, could be identified to the species level by eDNA (Figure 2). Damselfish, *Chromis chromis*, made up 46.3% of the visual observations at Dafni beach and 8.4% of UVC at Korakonissi but was not detected at species-level by eDNA, while the genus *Chromis* was detected at Dafni Beach.

UVC and eDNA both detected 13 genera (34 %), but at the species-level they shared only eight observations or 20 % of species-level detections (Figure 3, Supplementary Table 3), emphasizing the complementary nature of different survey methods. eDNA detected six more non-indigenous species than UVC, such as the Red Sea hardyhead silverside (*Atherinomorus forskalii*), a small Lessepsian species that would be difficult to tell apart from native sand smelts using visual census. Other non-indigenous species detected by eDNA included the Indo-Pacific red-toothed triggerfish (*Odonus niger*), Fangtooth moray (*Enchelycore anatine*), Hound needlefish (*Tylosurus crocodilus*), the marbled spinefoot (*Siganus rivulatus*) and the Bluespotted cornetfish (*Fistularia commersonii*). Ornate wrasse (*Thalassoma pavo*) was a considerable component of the community detected by both methods, making up 22.3% and 25.5% of eDNA detections and 10.7% and 22.8% of UVC observations, from Dafni Beach and Korakonissi, respectively (Figure 2). Parrotfish (*Sparisoma cretense*), dusky spinefoot (*Siganus luridus*), white seabream (*Diplodus sargus*), and salema porgy (*Sarpa salpa*) were also detected at both locations with both methods (Figure 2, Figure 3).

**Figure 3.**
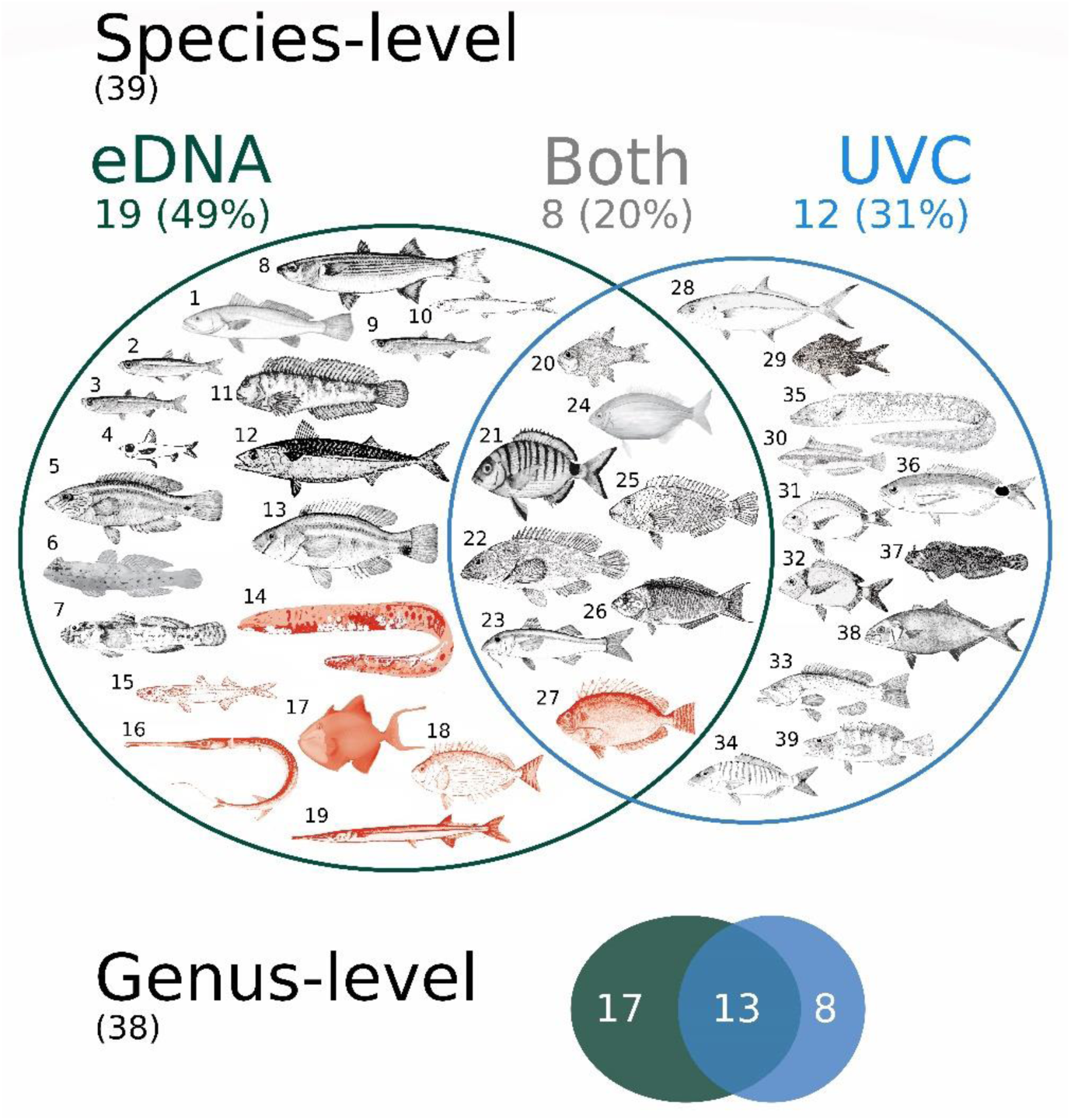
Venn diagram illustrating species-level detections and observations from aqueous eDNA and UVC respectively. Fish images and left aligned numbers correspond to the following species: 1) *Argyrosomus regius*, 2) *Atherina boyeri*, 3) *Atherina hepsetus*, 4) *Chlorophthalmus agassizi*, 5) *Symphodus melops*, 6) *Gobius bucchichi*, 7) *Gobius paganellus*, 8) *Oedalechilus labeo*, 9) *Sardina pilchardus*, 10) *Sardinella aurita*, 11) *Scartella cristata*, 12) *Scomber scombrus*, 13) *Symphodus cinereus*, 14) *Enchelycore anatina*, 15) *Atherinomorus forskalii*, 16) *Fistularia commersonii*, 17) *Odonus niger*, 18) *Siganus rivulatus*, 19) *Tylosurus crocodilus*, 20) *Apogon imberbis*, 21) *Diplodus sargus*, 22) *Epinephelus marginatus,* 23) *Mullus surmuletus*, 24) *Sarpa salpa*, 25) *Sparisoma cretense*, 26) *Thalassoma pavo*, 27) *Siganus luridus*, 28) *Caranx crysos*, 29) *Chromis chromis*, 30) *Coris julis*, 31) *Diplodus annularis*, 32) *Diplodus vulgaris*, 33) *Epinephelus costae*, 34) *Lithognathus mormyrus*, 35) *Muraena Helena*, 36) *Oblada melanura*, 37) *Parablennius gattorugine*, 38) *Seriola dumerili*, 39) *Serranus scriba*. Red fish images are non-indigenous species. The small Venn diagram provides the same data as the illustrated Venn diagram but shown at the genus-level and including detections or identifications that were specific to either species or genus. Additional metadata is in Supplementary Table 3.

### Sponge nsDNA as a molecular biomonitoring method

Four sponge species were biopsied, and their total DNA was extracted and prepared for vertebrate metabarcoding and Sanger sequencing (Table 1). Only *Chondrosia reniformis* was successfully amplified for COI identification, while the other three species were identified based on their skeletal (spongin) characteristics. *Chondrosia reniformis* (Figure 4A) occurred at both Dafni Beach and Korakonissi. *Sarcotragus* sp. (Figure 4B) and *Ircinia variabilis* (Figure 4C) were found only at Dafni Beach, while another *Ircinia* sp. (Figure 4D) was unique to Korakonissi. *Ircinia* sp. had the highest success rate, such that 44% of specimens collected resulted in over 100 reads with detections of the target taxa, Actinopterygii. When combining sponge samples, seven fish species were detected from nsDNA at Dafni Beach and 11 fish species were detected at Korakonissi (Figure 4E). Twenty-four unique detections were made at Dafni Beach when combining aqueous eDNA samples compared to seven from sponge samples. Of the seven sponge detections, *Mullus surmuletus* was the only fish not detected in the aqueous eDNA at Dafni Beach. Thirty-three unique detections were made at Korakonissi, while of the 11 detections from sponges, three were not present in the aqueous eDNA (i.e., *Gobius bucchichi*, *Gobius* sp., *Chromis* sp.). When comparing the total read counts from eDNA samples to the total reads of nsDNA, certain species were detected by the same order of magnitude (i.e., *Siganus luridus*, *Siganus rivulatus*, *Diplodus* sp., *Diplodus sargus* and *Thalassoma pavo*). Some cryptic species that could be found in closer proximity to sponges had more combined total reads in the nsDNA than eDNA, such as two goby species at Korakonissi (i.e. *Gobius bucchichi* and *Gobius paganellus*).

**Figure 4.**
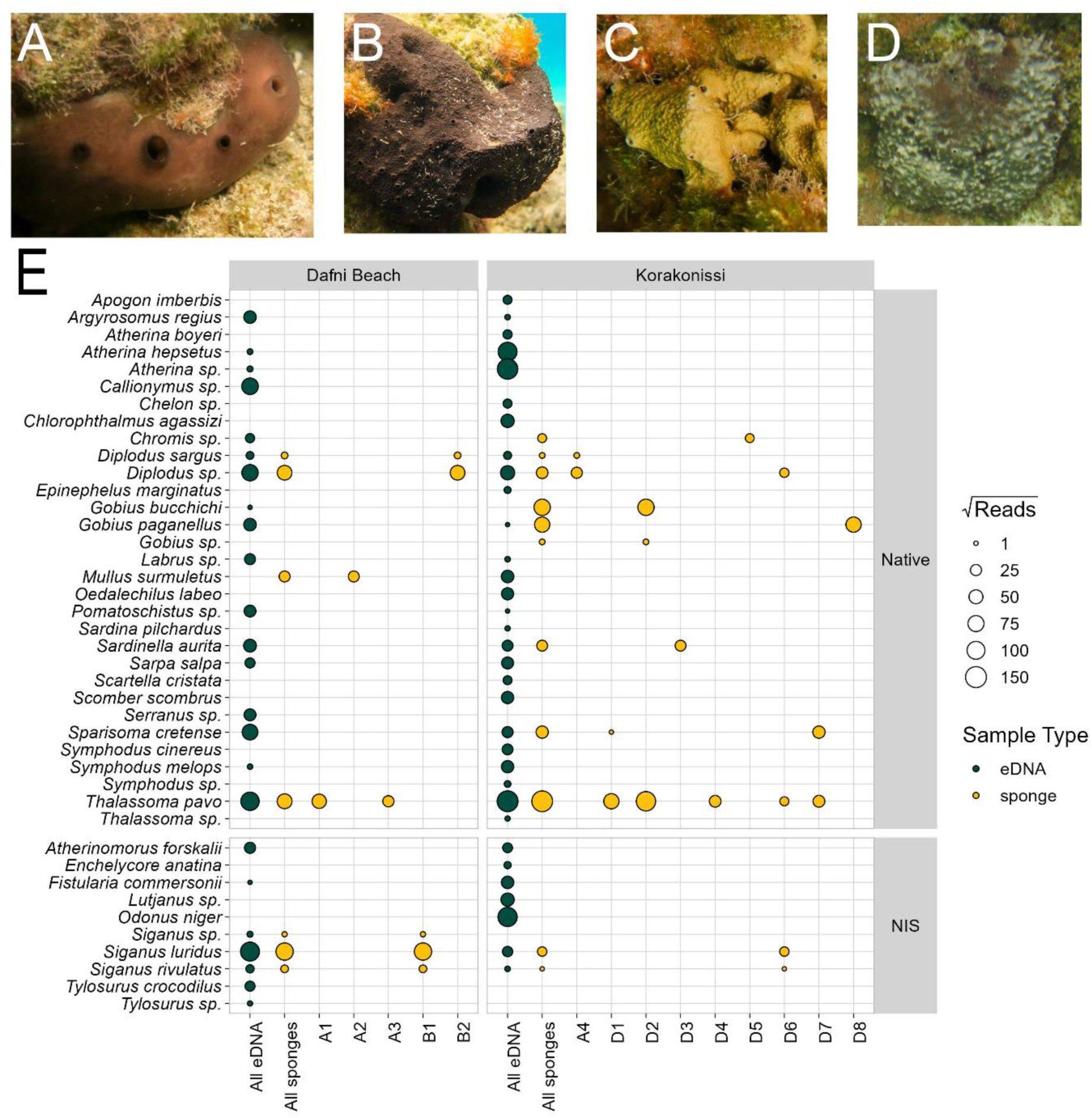
Pictures of the sponge species that were biopsied for sponge nsDNA analysis. **A)** *Chondrosia reniformis* **B)** *Sarcotragus* sp. **C)** *Ircinia variabilis* **D)** *Ircinia* sp. **E)** Bubble plot showing the square root reads of all eDNA and all sponge nsDNA samples combined as well as individual sponge nsDNA samples (sample letter corresponds to species A-D and sample number corresponds to replicate).

### Zakynthos, Greece marine island biodiversity

The Shannon Index at both Dafni Beach and Korakonissi did not significantly differ between eDNA and UVC sampling methods (Wilcoxon rank-sum: U = 17, p = 0.4306; U = 21, p = 0.0832; respectively). At Dafni Beach, the Shannon Index from sponges could only be calculated for two samples since species richness was low. However, at Korakonissi the sponges could be compared to the other sampling methods and were significantly different from UVC (Wilcoxon rank-sum: U = 21, p = 0.0004) but not eDNA (Wilcoxon rank-sum: U = 21, p = 0.0832) (Figure 5A). The J-Evenness at Dafni Beach for eDNA and UVC did not significantly differ (Wilcoxon rank-sum: U = 19, p = 0.5853). Similarly, at Korakonissi J-Evenness did not significantly differ between all pairwise combinations of sampling methods (Wilcoxon rank-sum: p > 0.05) (Figure 5B). The group dispersions by sampling location of UVC and eDNA beta-diversity were both homogenous (ANOVA: df = 1, F = 1.468, p = 0.254; df = 1, F = 3.412, p = 0.114, respectively). Beta-diversity (Bray-Curtis dissimilarity) significantly differed between locations in both the UVC data and eDNA data (PERMANOVA: df = 1, R^2^ = 0.60, F = 15.088, p = 0.004; df = 1, R^2^ = 0.33, F = 2.937, p = 0.027; respectively). The results remained the same when using the Jaccard index for UVC data but were no longer significant for eDNA (PERMANOVA: df = 1, R^2^ = 0.320, F = 4.701, p = 0.003; df = 1, R^2^ = 0.221, F = 1.705, p = 085; respectively).

**Figure 5.**
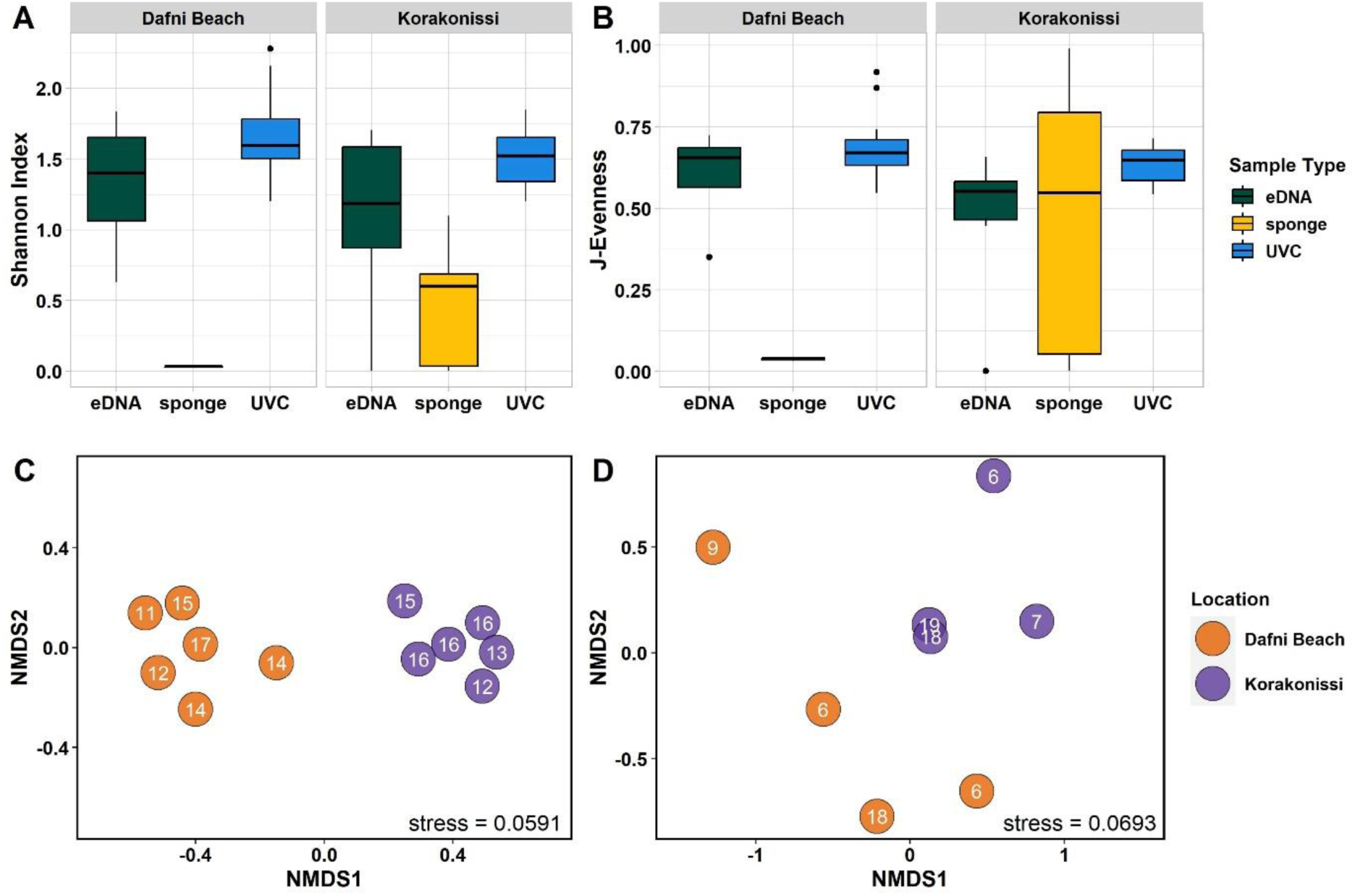
A) Box plots of Shannon indices calculated for the different sampling types at each location. **B)** Box plots of J-evenness calculated for the different sampling types at each location. **C)** NMDS plot calculated from Bray-Curtis dissimilarity calculated from UVC, where numbers on points indicate alpha-diversity. **D)** NMDS plot calculated from Bray-Curtis dissimilarity from eDNA, where numbers on points indicate alpha-diversity.

The aqueous eDNA samples from this study were compared to a previous study on eDNA sampled from inside and outside the National Marine Park of Zakynthos (Aglieri et al., 2021) (Figure 6). The eDNA samples from these different studies and sites had similar species accumulation curves, suggesting a similar number of fish species expected in the locations. One exception to this observation were samples taken from the MPA at the previous study, which have about half the number of species compared to our samples, despite a similar sample size.

**Figure 6.**
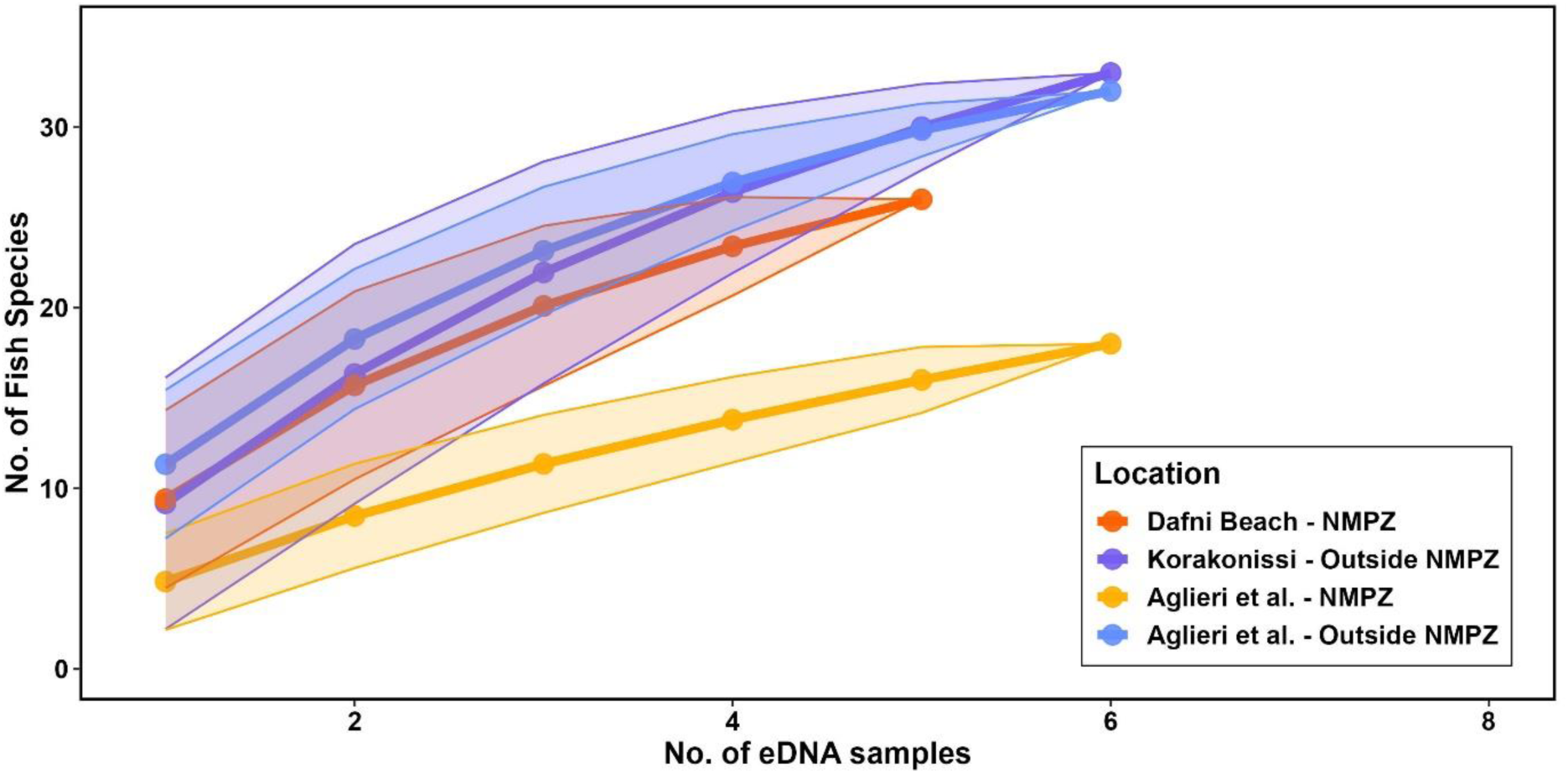
Species accumulation curves of aqueous eDNA samples from this study and from Aglieri and colleagues 2021 inside and outside the MPA of the National Marine Park of Zakynthos (NMPZ).

## Discussion

Monitoring biological elements within MPAs over the long term poses a significant challenge to managers and conservation practitioners in terms of effort, cost, capacity, and efficiency. Timely and precise information on the status of protected elements (i.e., species, habitats) is critical for their effective management, particularly when considering how climate change is calling for more adaptive and climate smart management strategies. Environmental DNA could be a solution to obtain a more comprehensive inventory of marine biodiversity information especially in MPAs, while potentially acting as an early warning system for non-indigenous species for which implementing counteracting measures at early invasion stages is crucial for a successful management outcome (Simberloff et al., 2013). Sponge specimen collection for natural sampler DNA (nsDNA) could be promising for establishing baselines, where specimens have been collected and stored prior to the advent of eDNA technology. The present study on fish diversity in Zakynthos offers important insights into the benefits and caveats of implementing monitoring strategies using complementary methods.

### Aqueous eDNA Detections vs. UVC observations

Like previous research studies, eDNA was able to provide greater taxonomic granularity than UVC and both methods provided complementary information (Aglieri et al., 2021; Lamy et al., 2021; Valdivia-Carrillo et al., 2021). For instance, at both locations the wrasse genus *Symphodus*, was recorded by UVC, but the species *Symphodus cinereus* and *S. melops* were detected only by eDNA (Figure 2). Often this lack of species-level identification visually, was what led to a low number of species (21%) present in both data sets, whereas genus-level overlap between the data sets was much greater (37%) (Figure 3). Yet, eDNA was capable of detecting the presence of fish species inhabiting deep waters as adults and shallow pelagic areas as larvae and juveniles (e.g. *Chlorophthalmus agassizi*)(D’Onghia et al., 2006) that inevitably go unnoticed by UVC methods due to depth limitation and difficulty of visually spotting and identifying species at early life stages. There were exceptions where UVC distinguished more species within a genus than eDNA metabarcoding. While both UVC and eDNA detected the white seabream *Diplodus sargus*, eDNA failed to identify *D. annularis* and *D. vulgaris*, which were instead visually detected. Similarly, *Chromis chromis* was abundant at both locations, however only genus-level (*Chromis* sp.) detections were present in the eDNA from Dafni Beach and the sponge nsDNA from Korakinissi. This ambiguity in the molecular methods, despite obvious visual presence, was due to the inability of the primer we used to resolve the *Chromis* genus, an occasional trade-off of metabarcoding analysis (Zhang et al., 2018). Although only one *Chromis* species is endemic to the Mediterranean, further damselfish species within this genus could expand their ranges into the Mediterranean and potentially go undetected by this method. This further highlights the importance of treating primers as assays in the sense that even though they are designed to universally amplify fish taxa, there will always be closely-related species that may remain indistinguishable, while others will have sequences with poor affinity with the chosen primers. Despite these minor ambiguities – and the impact of the aggressive extraction protocol – the eDNA metabarcoding analysis still produced a more comprehensive list of species than did the UVC survey (Figure 3).

When considering individual samples, fish diversity (Shannon index) and evenness detected by eDNA and UVC did not significantly differ (Figure 5A, 5B) and both methods were able to distinguish location specific beta-diversity (Figure 5C, 5D). Environmental DNA metabarcoding has been used to determine beta-diversity of fish communities on spatial scales relevant to management, and at a regional scale it appears to detect more taxa than UVC (Lamy et al., 2021), which is similar to what we observed in Zakynthos. In tropical environments with high diversity, eDNA metabarcoding is often able to capture a wider variety of fish families than UVC (Polanco Fernández et al., 2021; Zamani et al., 2022) and we also observe this, with eDNA capturing nine (36% of total) unique families, two of which include NIS.

### Detection of Non-indigenous species

eDNA and sponge nsDNA showed greater detection ability of non-indigenous species relative to UVC. eDNA metabarcoding detected seven NIS (i.e. *Atherinomorus forskalii*, *Enchelycore anatine, Fistularia commersonii*, *Odonus niger*, *Siganus luridus, Siganus rivulatus*, *Tylosurus crocodilus*), while only one NIS (*Siganus luridus*) was observed with UVC (Figure 2). Although the fish fauna in Zakynthos protected areas have been systematically and regularly monitored over the last decade by the use of UVC, fisheries catch data and citizen science initiatives with a special emphasis on detecting NIS (Dimitriadis et al., 2021, 2024; Dimitriadis, et al., 2023; Ragkousis et al., 2023). Red Sea hardyhead silverside (*Atherinomorus forskalii*), red-toothed triggerfish (*Odonus niger*), and houndfish (*Tylosurus crocodilus*) to our knowledge have not been previously reported in our study area. However, *Atherinomorus forskalii* is not new to Greek waters (Bariche et al., 2015; Zenetos et al., 2013) and was reported in Zakynthos in 2022 on a citizen science platform (i.e., iNaturalist; https://greece.inaturalist.org/taxa/608753-Atherinomorus-forskalii), emphasising the power of citizen science. The eDNA detection of red-toothed triggerfish (*Odonus niger*), to the best of our knowledge, is the first record of this species in the Mediterranean region. While *Odonus niger* is native to the Red Sea it is also an ornamental fish species sold in Greek aquarium stores for direct trade (Papavlasopoulou et al., 2014), rendering accidental or deliberate release by aquarists as a possible mode of introduction in the study area. Houndfish (*Tylosurus crocodilus*) was first reported in the North Aegean Sea in 2005 (Sinis, 2005) and more recently in the Adriatic Sea (Fortič et al., 2023), such that the present finding is the third report of the species in the Mediterranean.

The most established non-indigenous taxa in Zakynthos are the rabbitfishes and their herbivory is causing shifts in ecosystem function (Dimitriadis et al., 2021; Golani, 1998). Dusky spinefoot (*Siganus luridus*) was introduced to Zakynthos in 2004 and today comprises the most abundant fish species in the National Marine Park of Zakynthos (NMPZ); it accounts for almost one third of total fish biomass (Dimitriadis et al., 2024) which interestingly mirrored the relative read abundance recorded in this study (26.4%) at Dafni Beach. Marbled spinefoot (*Siganus rivulatus*) was observed for the first time in NMPZ in 2009 (Dimitriadis et al., 2021) and progressively increased its population size (Dimitriadis et al., 2024). Both rabbitfish species were present in the sponge nsDNA from both Dafni Beach and Korakonissi.

Snappers (genus: *Lutjanus*) are known Lessepsian migrants, which we detected at Korakonissi and species from this genus are known to have colonised the Mediterranean since the 1970s (Bariche et al., 2015). Moreover, mangrove red snapper (*Lutjanus argentimaculatus*) was recently detected by eDNA metabarcoding in the NMPZ (Aglieri et al., 2021). While the physical ability of these fish to enter via the Suez Canal has facilitated the Mediterranean to become a NIS hotspot, ocean warming of the region is what ultimately allows these warm-affinity fish to thrive. With eDNA we detected the range expanding fangtooth moray (*Enchelycore anatina*), a subtropical species native to the eastern Atlantic, which since it was first reported in the study area in 2012 (Kapiris et al., 2014) it has been rare to appear in fish monitoring surveys.

### Refining the Molecular Tool kit

eDNA metabarcoding analysis has been explored as a method for early and rapid detection of NIS (Duprey et al., 2023; Zaiko et al., 2015). But a more common approach when studying NIS with eDNA is to use species-specific primers with analyses such as PCR (Chucholl et al., 2021), qPCR (LeBlanc et al., 2020) and ddPCR (Chucholl et al., 2021; Doi et al., 2015; Nathan et al., 2014). When comparing species detection using species-specific approaches with metabarcoding, the latter has been able to perform similarly to qPCR and for the same investigator effort delivers multiple species distribution data (Harper et al., 2018). Of course, the outcome of these methods comparisons could differ depending on the species of interest. Metabarcoding analysis is useful in exploratory contexts such as our study, where many of the invasive species detected were not yet observed by UVC in the study area, so they were relatively unexpected.

Sponge nsDNA is an emerging tool for biodiversity census in aquatic systems (Gallego et al., 2024; Mariani et al., 2019; Turon et al., 2020), and this study provides valuable information on how different sponge species function as natural environmental DNA samplers. In previous studies, captive sponges have been shown to vary in eDNA signal based on species (Cai et al., 2022) and evidence of this in the natural world also exists (Brodnicke et al., 2023; Neave et al., 2023), despite some conflicting observations showing no differences between sponge species, but this could have been due to low replication (Turon et al., 2020). Sponges have different pumping rates which can change based on environmental conditions and contain different levels of microbial symbionts that could inhibit the DNA extraction processes (Brodnicke et al., 2023.; Harper et al., 2023). In particular, our targeted sponge species are notoriously abundant in microbial symbiotic partners, which usually leads to poor filtration rates (Weisz et al., 2008). Only a few studies have compared eDNA signal from sponge nsDNA to that of aqueous eDNA. The results are varied but remarkable, a couple findings highlighting stark contrasts between sponge nsDNA and aqueous eDNA (Brodnicke et al., 2023; Jeunen et al., 2023), while another found no significant difference between the methods (Jeunen et al., 2021). Further, a recent study suggests that sponge nsDNA may detect more complex community structures compared to aqueous eDNA (Cai et al. 2024). Clearly the sponge species, laboratory processes and environmental factors all could be playing a role in the efficiency of these natural samplers. All of the sponges sampled in this study were relatively ineffective compared to previous efforts using sponge nsDNA for biodiversity surveys (Jeunen et al., 2021; Neave, et al., 2023; Riesgo et al., 2024), but were comparable to some efforts where individual samples consistently detected less than ten species (Brodnicke et al., 2023; Turon et al., 2020). Based on our results, it appears that further research is necessary to thoroughly understand what determines the suitability of sponge species for nsDNA analyses. In previous studies, *Chondrosia reniformis* was also tested and did not have any detections of target taxa (Mariani et al., 2019) but without comparison to water and based on only one sample, it was not certain whether the lack of detections was representative of this species. Despite the suboptimal performance of the sponges in this study, our nsDNA analysis still led to the detection of two non-indigenous species and 13 taxa in total.

Tropicalization involves the range expansions of tropical species and retractions of native, more temperate species. Prior studies have used eDNA to track range expansions of invasive (Larson et al., 2017; Richardson et al., 2016) and endangered species (Harper et al., 2019; Hobbs et al., 2019; Valsecchi et al., 2022). However, many of these studies focus on a single species and have developed species-specific probes, which is why the ability for eDNA to monitor tropicalization needs further study using metabarcoding analysis to monitor the expansion and contraction of native and non-native species. It is interesting to note that the species-specific molecular strategies seem to mirror long-standing traditional conventions of environmental management, which initially placed much emphasis on single species, and only very recently are beginning to move to a more holistic approach. Species-specific probes may be more attractive to management because they may provide a sense of a more controlled, bespoke survey, but in terms of monitoring tropicalization, metabarcoding analysis has more potential. Whilst metabarcoding analysis can lead to false positives (Darling et al., 2021) and false negatives (Jackman et al., 2021), there are methodological strategies to overcome these challenges (McClenaghan et al., 2020) and overall it provides multiple species distributions simultaneously (Harper et al., 2018) and may be more effective at early detection of non-indigenous fishes (Van Nynatten et al., 2023). While the presence of tropical NIS is evidence of topicalization, long term molecular monitoring coupled with other abiotic measurements could lead to further insights regarding the rate at which this is occurring, and its ecosystem ripple effects. One such study over a longer temporal scale used eDNA metabarcoding of archived samples to show short-term tropicalization of fish communities over a marine heatwave (Gold et al., 2023). Our study presents seven species-level and three genus-level detections of tropical NIS, only two of which (*Lutjanus* sp. and *Siganus luridus*) were detected in eDNA samples collected from the NMPZ only three (i.e. June, July 2018) years prior (Aglieri et al., 2021). This difference highlights that repeat molecular monitoring could lead to insights on tropicalization that UVC does not have the sensitivity for, which would represent the early warning system that MPA managers require for the development of timely and effective NIS mitigation strategies.

Environmental DNA and UVC were both able to distinguish Mediterranean fish communities, from high and low protection status MPAs, less than 30 km apart. Exemplifying the complementary nature of eDNA and UVC, community composition at each location differed such that only seven (13%) and 10 (18%) of 55 taxa were detected by both across locations which corroborates many prior studies showing that eDNA and UVC typically detect different sets of species (Aglieri et al., 2021). Still eDNA provided nearly half (49%) of the species detections alone and captured six out of seven NIS, from which three are new records in the study area despite the regular, long-standing and NIS focused fish monitoring at Zakynthos islands by visual census and fisheries related methods (Dimitriadis et al., 2023). Sponge nsDNA captured 13 taxa, all of which were also detected by eDNA. Despite likely suffering from PCR inhibition, sponge nsDNA successfully detected dominant species including sea breams (*Diplodus* sp.), non-indigenous rabbit fishes (*Siganus* sp.) and ornate wrasses (*Thalassoma pavo*). *Siganus luridus* was the only NIS detected by all three methods, highlighting the strength of molecular tools for determining NIS presence. The red-toothed triggerfish (*Odonus niger*), an Indo-Pacific species present in the Red Sea, was detected at Korakonissi and to our knowledge has never been previously reported in the Mediterranean, while *Tylosurus crocodilus* is only the third record in this region. Overall, this study advocates for a more integrated approach for marine biodiversity monitoring, combining UVC with eDNA metabarcoding. Such a holistic approach is crucial for early detection of ecosystem changes, particularly in the context of increasing tropicalization and the introduction of NIS, enabling more proactive and informed conservation strategies. This approach also offers a comprehensive understanding of marine ecosystems, ensuring a balance between scientific rigor and practical applicability in biodiversity monitoring and management.

## Acknowledgments

We would like to thank the staff responsible for helping us with the logistics and paperwork associated with sampling in the National Marine Park of Zakynthos. We also are grateful for assistance with sponge identification by Sanger sequencing and morphology. We thank Lynsey Harper for discussions on the field sampling plan. The study was supported by Grant NE/T007028/1 (SpongeDNA) from the UK Natural Environment Research Council to S.M. and A.R. A.R. was also supported by grants SponBIODIV (a 2021-2022 BiodivProtect joint call for research proposals, under the Biodiversa+ Partnership co-funded by the European Commission, and with the funding organisations ‘Fundación Biodiversidad’ and FORMAS), a Consolidación Investigadora CNS2023-144571 from MICIU/AEI/10.13039/501100011033/ and “Unión Europea NextGenerationEU/PRTR” funds, and an intramural grant from CSIC (PIE-202030E006).

## Notes

https://zenodo.org/records/15392014?token=eyJhbGciOiJIUzUxMiJ9.eyJpZCI6IjIxMDIyMzgyLTUxZDktNDM2MS1hMzEwLTdhNmUwYjFiMTIyMSIsImRhdGEiOnt9LCJyYW5kb20iOiJmMDEyMjA1MDYyNzhkNjVjNzZjYjU0YmUzM2NlNzUyZiJ9.HpGj3LzAMPPALaGWA0Sy08Uiq7LOliNS6MfZeU6W3lr5VmNC_G_kcNwk3PdkZl33FuMz4tnJ_yQRR7dc0xkpVQ

### Competing Interest Statement

The authors have declared no competing interest.

