## supplementary for "Refining a molecular tool kit to capture tropicalization in Mediterranean Marine Protected Areas"

*Underwater Visual Census*

Underwater visual census (UVC) was performed within a ~10 meter sampling radius and using the standard stationary point methods established by the U.S. National Marine Fisheries Service (Bohnsack & Bannerot, 1986). UVC was achieved with two snorkelling observers, each with over 25 years of experience with identifying Mediterranean fish species. For 10 minutes per census, fish were identified to the lowest taxonomic rank possible and counted. However, schooling fish ≥~50 were recorded as an exact abundance of 50.

*Field Sampling Controls*

Filtration blanks and field blanks were collected to control for any contamination that may have occurred during the field sampling. All blanks were collected by filtering 2L of bottled drinking water. The filtration blanks were collected at the beginning and end of each sampling day (September 10-14, 2021). This resulted in eight filtration blanks, the purpose of which was to check if there was any contamination in any filtration equipment such as gloves, bleach bottles, paper towels, etc. The field blanks were taken at each sampling location change (where each sampling location was visited at two separate times, so sites could be resampled). This resulted in four field blanks, the purpose of which was to have an extra level of precaution, so if contamination appeared, we'd have the best chance of identifying when and where it might have happened.

*Sponge Identification*

Sponges are notoriously difficult to identify based on morphology alone, so a fragment of the *cytochrome c oxidase I* (COI) was used to identify and confirm species. Each sponge sample was PCR amplified using the universal primers known for their efficiency in amplifying invertebrate DNA, using LCO1490 (5'-GGTCAACAAATCATAAAGATATTGG-3') and matched to HCO2198 (5'-

TAAACTTCAGGGTGACCA AAAAATCA-3’) (Folmer et al. 1994), to amplify a 658 bp fragment of the COI. The PCR protocol included a 10 min denaturing step at 95°C, followed by 35 cycles of 95°C for 1 min, 48-52°C for 1 min, and 72°C for 30 sec, with a final extension at 72°C for 5 min. Amplified DNA was Sanger sequenced in both directions at Macrogen.

*Sequencing*

The aqueous eDNA library was pooled at equimolar concentration with a library from another project for sequencing, while the sponge nsDNA library was sequenced on a dedicated separate run. The final libraries were diluted to a molarity of 85pM with a 10% PhiX spike-in and sequenced at Liverpool John Moores University on separate runs (aqueous eDNA - October 2022; sponge nsDNA – December 2022) using an Illumina iSeq100 with iSeq i1 Reagent v2 300 cycles. The eDNA sequencing run resulted in 7,749,842 reads total, of which 4,099,666 reads pertained to this project (the aqueous eDNA library was sequenced alongside a separate project), and the sponge sequencing run resulted in 7,052,976 reads total.

*Bioinformatics Pipeline*

The sequence libraries were analyzed using OBITools 1.2.11 (Python v2) (Boyer et al., 2016). Raw R1 and R2 sequences were trimmed to a length of 149 bp using the command *obicut* to remove bases determined from the per-base quality scores from *fastqc*. The trimmed reads were merged using *illuminapairedend*, keeping paired-end alignments at >30 quality score. Paired-end alignments were then demultiplexed using *ngsfilter* and the identified barcoded samples (aqueous eDNA and sponge DNA) were concatenated into a single fasta file. The sequences were then filtered by length (130 - 190 bp) and dereplicated using *obiuniq*. Chimeras were removed *de novo* using VSEARCH version 2.4.3 (Rognes et al., 2016). The remaining sequences were then clustered using SWARM v2 (Mahé et al., 2015) with d value = 1.

Taxonomy was assigned by searching for consensus between two methods: 1) *ecotag* with a database created using *ecoPCR* with the Tele02 primers querying the EMBL database (release version r143) for non-human vertebrates (taxid settings: -r 7742 -i 9606) and 2) Bayesian LCA-based taxonomic classification method (BLCA) (Gao et al., 2017) with a custom 12S reference database, generated using a custom python script: https://github.com/shump2/Haploconator/blob/main/python_scripts/0.generate_dbBLCA.py. Briefly, accession numbers matching to the mitochondrial 12S ribosomal RNA were downloaded from Genbank and the function *blastdbcmd* was used to extract sequences from the nt_euk and ref_euk NCBI databases (accessed June 2022). A blast database was generated using *makeblastdb* and the taxonomy file used for BLCA was formatted. The taxonomic output was merged by the molecular operational unit (MOTU) IDs. If the taxonomies matched, the assignment with the highest percent identity was used, but if the taxonomies differed (e.g. *ecotag* assigned to family-level while BLCA assigned to genus-level), the taxonomy with the highest percent identity > 90% and at the most informative taxonomic level was retained (e.g. if the family-level assignment has a percent identity of 100% and the genus-level has a percent identity of 98%, the detection was assigned to the genus-level taxonomy). Human reads were removed. Both assignment methods together (Supplementary Table 2) supplemented by manual checks using the NCBI BLAST tool (Supplementary Table 4), resulted in the best taxonomic resolution of the dataset. Taxonomies were collapsed using a 98% similarity method whereby species and genus-level assignments were retained if they had ≥ 98% identity, family-level assignments were retained if they had 90-100% identity, order-level assignments were retained if they had 80-100% identity, and class and phylum level assignments were retained if they had 70-100% identity. Species and genus-level assignments were only retained if they were ≥ 98% to have high confidence in the taxonomy assigned, particularly in the context of non-indigenous species identification. This ≥ 98% taxonomic assignment and removal of human reads resulted in 324,050 reads or ~14,700 reads per sample.

### *Supplementary References*

Boyer, F., Mercier, C., Bonin, A., Le Bras, Y., Taberlet, P., & Coissac, E. (2016). obitools: A unix-inspired software package for DNA metabarcoding. *Molecular Ecology Resources*, *16*(1), 176–182. https://doi.org/10.1111/1755-0998.12428

Folmer, O., Black, M., Hoeh, W., Lutz, R., & Vrijenhoek, R. (1994). DNA primers for amplification of mitochondrial cytochrome c oxidase subunit I from diverse metazoan invertebrates. *Molecular Marine Biology and Biotechnology*, *3*(5), 294–299.

Gao, X., Lin, H., Revanna, K., & Dong, Q. (2017). A Bayesian taxonomic classification method for 16S rRNA gene sequences with improved species-level accuracy. *BMC Bioinformatics*, *18*(1), 247. https://doi.org/10.1186/s12859-017-1670-4

Mahé, F., Rognes, T., Quince, C., de Vargas, C., & Dunthorn, M. (2015). Swarm v2: Highly-scalable and high-resolution amplicon clustering. *PeerJ*, *3*, e1420. https://doi.org/10.7717/peerj.1420

Rognes, T., Flouri, T., Nichols, B., Quince, C., & Mahé, F. (2016). VSEARCH: A versatile open source tool for metagenomics. *PeerJ*, *4*, e2584. https://doi.org/10.7717/peerj.2584

### Supplementary Tables

**Supplementary Table 1.** Contamination found in controls and the corresponding number of reads which were removed from the dataset.

| Contaminant | Found In | Reads | Reads removed from Dataset |
|---|---|---|---|
| *Atherina hepsetus* | sample.DBE_filterstart | 2 | 20 |
| *Atherina* sp. | sample.DBM_fieldblank | 1 | 10 |
| *Dicentrarchus labrax* | sample.pcrpositive4 | 1 | not present in other samples |
| *Gobius bucchichi* | sample.pcrpositive | 1 | 10 |
| *Lagodon rhomboides* | sample.pcrnegative4 | 42 | not present in other samples |
| *Odonus niger* | sample.DBE_filterend | 1 | 10 |
| *Scomber scombrus* | sample.DBE_filterstart | 2 | 20 |
| *Sprattus sprattus* | sample.pcrpositive4 | 1 | 10 |
| *Symphodus bailloni* | sample.pcrpositive4 | 1 | not present in other samples |
| *Thalassoma pavo* | sample.pcrnegative | 1 | 10 |
| *Thalassoma* sp. | sample.pcrnegative4 | 82 | 820 |

**Supplementary Table 2.** Left column showing MOTUs which were unassigned, assigned, assigned to target taxa and target taxa at genus or species level over the three steps of taxonomic assignment. Right column showing number of taxa for each taxonomic level after MOTUs are collapsed over the three steps of taxonomic assignment. The final dataset information is in row three, which combines all taxonomic assignment methods.

| | **MOTUs** | | | | **Collapsed Taxonomy** | | | | |
|---|---|---|---|---|---|---|---|---|---|
| | Unassigned MOTUs | Assigned MOTUs | Target Taxa MOTUs | Target Taxa MOTUs at Genus or Species Level | Species (100% - 98%) | Genus (100% - 98%) | Family (100% - 90%) | Order (100% - 80%) | Class and Phylum (100% - 70%) |
| **1. ecotag w/ EMBL database of target taxa only** | 537 | 535 | 535 | 285 | 23 | 20 | 42 | 6 | 49 |
| **2. BLCA w/ custom reference database** | 560 | 512 | 238 | 217 | 18 | 31 | 33 | 4 | 2 |
| **3. Combined methods w/ blast** | 269 | 803 | 536 | 417 | 27 | 14 | 8 | 3 | 6 |

**Supplementary Table 3.** Species and genera detections displayed in Figure 3.

| Label # | Scientific name | Venn |
|---|---|---|
| 1 | *Argyrosomus regius* | eDNA |
| 2 | *Atherina boyeri* | eDNA |
| 3 | *Atherina hepsetus* | eDNA |
| 4 | *Chlorophthalmus agassizi* | eDNA |
| 5 | *Symphodus melops* | eDNA |
| 6 | *Gobius bucchichi* | eDNA |
| 7 | *Gobius paganellus* | eDNA |
| 8 | *Oedalechilus labeo* | eDNA |
| 9 | *Sardina pilchardus* | eDNA |
| 10 | *Sardinella aurita* | eDNA |
| 11 | *Scartella cristata* | eDNA |
| 12 | *Scomber scombrus* | eDNA |
| 13 | *Symphodus cinereus* | eDNA |
| 14 | *Enchelycore anatina* | eDNA |
| 15 | *Atherinomorus forskalii* | eDNA |
| 16 | *Fistularia commersonii* | eDNA |
| 17 | *Odonus niger* | eDNA |
| 18 | *Siganus rivulatus* | eDNA |
| 19 | *Tylosurus crocodilus* | eDNA |
| 20 | *Apogon imberbis* | Both |
| 21 | *Diplodus sargus* | Both |
| 22 | *Epinephelus marginatus* | Both |
| 23 | *Mullus surmuletus* | Both |
| 24 | *Sarpa salpa* | Both |
| 25 | *Sparisoma cretense* | Both |
| 26 | *Thalassoma pavo* | Both |
| 27 | *Siganus luridus* | Both |
| 28 | *Caranx crysos* | UVC |
| 29 | *Chromis chromis* | UVC |
| 30 | *Coris julis* | UVC |
| 31 | *Diplodus annularis* | UVC |
| 32 | *Diplodus vulgaris* | UVC |
| 33 | *Epinephelus costae* | UVC |
| 34 | *Lithognathus mormyrus* | UVC |
| 35 | *Muraena helena* | UVC |
| 36 | *Oblada melanura* | UVC |
| 37 | *Parablennius gattorugine* | UVC |
| 38 | *Seriola dumerili* | UVC |
| 39 | *Serranus scriba* | UVC |

| # | Genus | Venn |
|---|---|---|
| 1 | *Argyrosomus* | eDNA |
| 2 | *Atherinomorus* | eDNA |
| 3 | *Callionymus* | eDNA |

| | | |
|---|---|---|
| 4 | *Chelon* | eDNA |
| 5 | *Chlorophthalmus* | eDNA |
| 6 | *Enchelycore* | eDNA |
| 7 | *Fistularia* | eDNA |
| 8 | *Labrus* | eDNA |
| 9 | *Lutjanus* | eDNA |
| 10 | *Odonus* | eDNA |
| 11 | *Oedalechilus* | eDNA |
| 12 | *Pomatoschistus* | eDNA |
| 13 | *Sardina* | eDNA |
| 14 | *Sardinella* | eDNA |
| 15 | *Scartella* | eDNA |
| 16 | *Scomber* | eDNA |
| 17 | *Tylosurus* | eDNA |
| 18 | *Apogon* | Both |
| 19 | *Atherina* | Both |
| 20 | *Chromis* | Both |
| 21 | *Diplodus* | Both |
| 22 | *Epinephelus* | Both |
| 23 | *Gobius* | Both |
| 24 | *Mullus* | Both |
| 25 | *Sarpa* | Both |
| 26 | *Serranus* | Both |
| 27 | *Siganus* | Both |
| 28 | *Sparisoma* | Both |
| 29 | *Symphodus* | Both |
| 30 | *Thalassoma* | Both |
| 31 | *Caranx* | UVC |
| 32 | *Coris* | UVC |
| 33 | *Lithognathus* | UVC |
| 34 | *Muraena* | UVC |
| 35 | *Oblada* | UVC |
| 36 | *Parablennius* | UVC |
| 37 | *Scorpaena* | UVC |
| 38 | *Seriola* | UVC |

**Supplementary Table 4.** MOTUs which were assigned to species level after using the NCBI Blast tool.

| MOTU id | best_identity_final | Manual Taxanomic assignment |
|---|---|---|
| gre2_000110514 | 100 | *Argyrosomus regius* |
| gre2_000111343 | 100 | *Atherinomorus forskalii* |
| gre2_000120754 | 100 | *Diplodus sargus* |
| gre2_000123570 | 100 | *Epinephelus marginatus* |
| gre2_000002161 | 99.4 | *Sardinella aurita* |
| gre2_000110418 | 99.26 | *Sarpa salpa* |
| gre2_000000080 | 100 | *Siganus luridus* |
| gre2_000013666 | 100 | *Siganus rivulatus* |
| gre2_000000578 | 100 | *Sparisoma cretense* |
| gre2_000110705 | 99.3939394 | *Symphodus cinereus* |
| gre2_000110841 | 99.39 | *Symphodus melops* |
| gre2_000000028 | 99.38 | *Thalassoma pavo* |
| gre2_000111411 | 99.38 | *Tylosurus crocodilus* |

**Supplementary Figures**

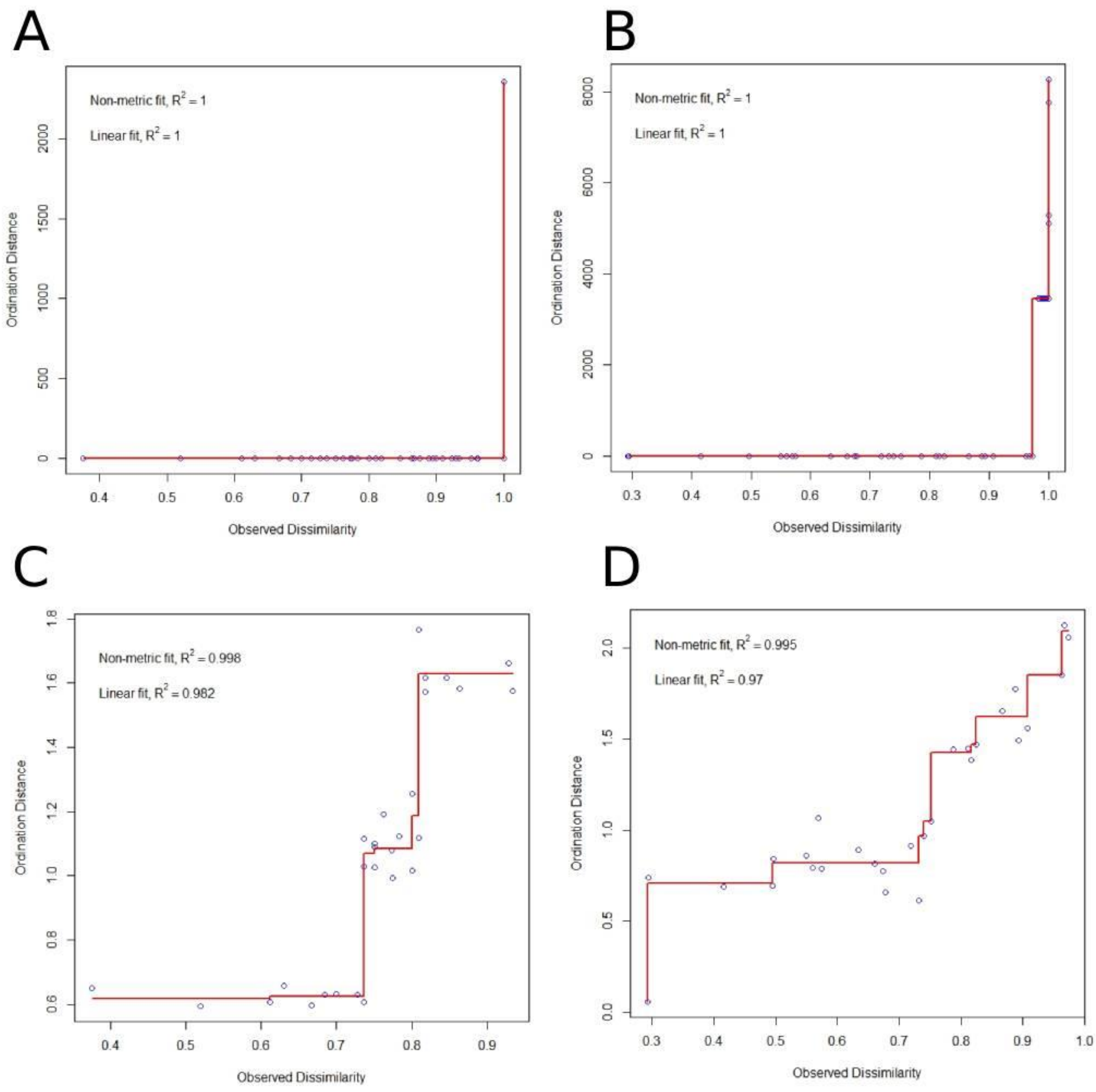


**Supplementary Figure 1.** Stress plots of eDNA ordination data. **A** Stress plot of Jaccard dissimilarity including all eDNA samples. **B** Stress plot of Bray-Curtis dissimilarity including all eDNA samples. **C** Stress plot of Jaccard dissimilarity with the three outlying samples are removed. **D** Stress plot of Bray-Curtis dissimilarity with the three outlying samples are removed.

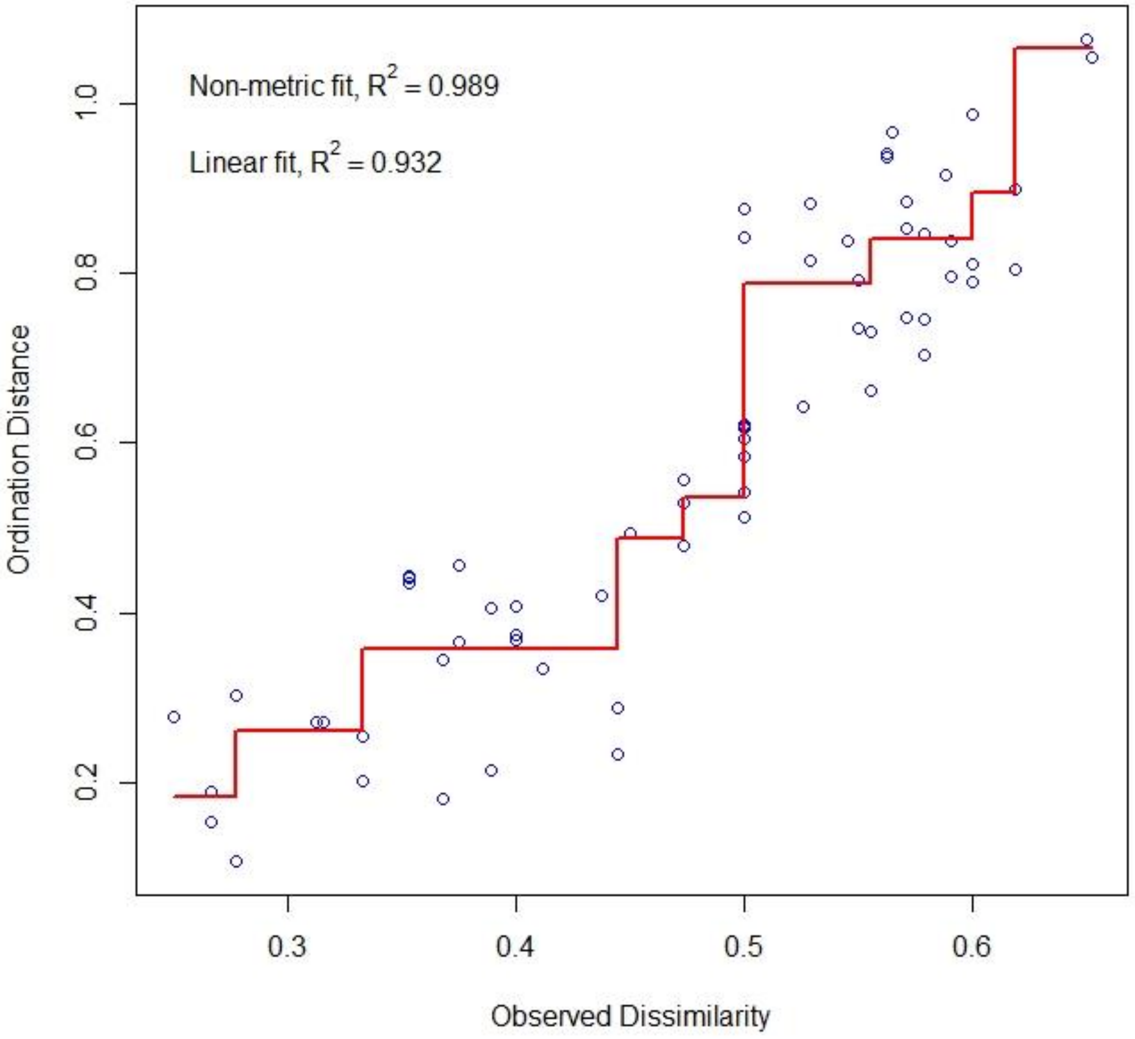


**Supplementary Figure 2.** Stress plot of Jaccard dissimilarity calculated from UVC data.

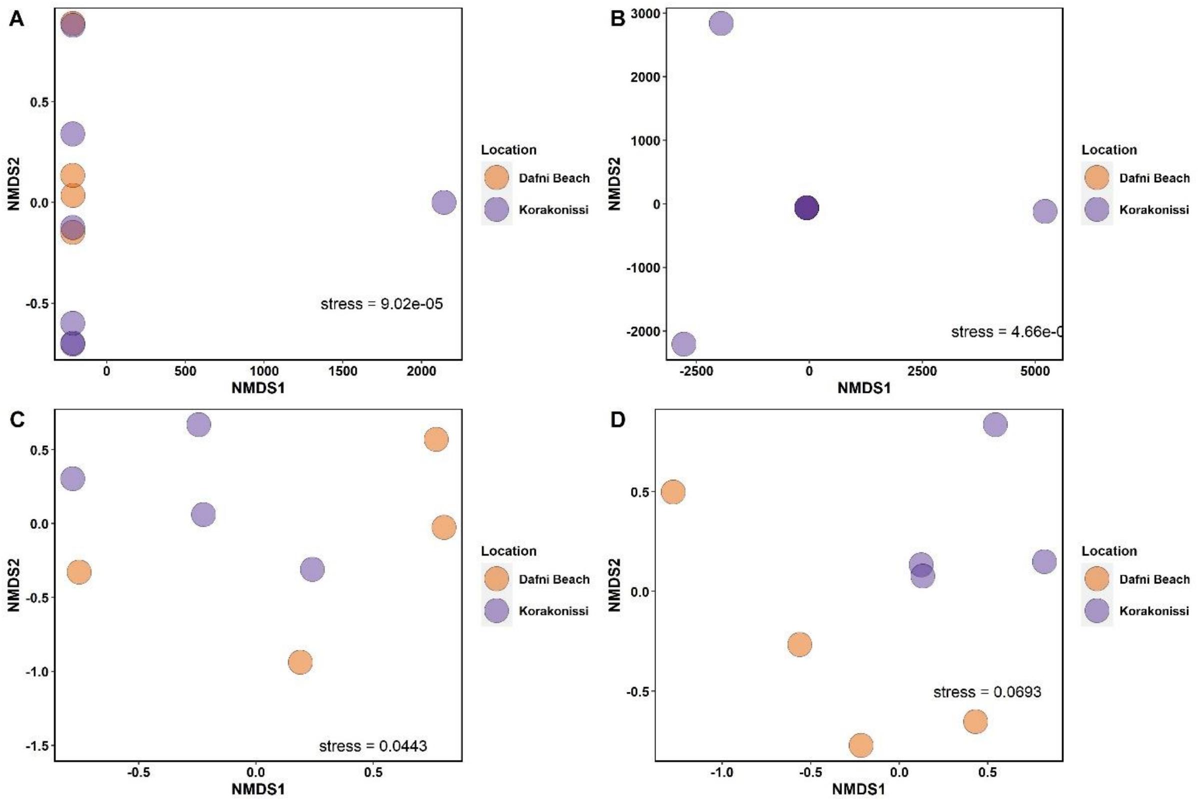


**Supplementary Figure 3**. NMDS plots of eDNA ordination data. All points are made slightly transparent, so darker shades (relative to legend) represent overlapping points. **A** NMDS of Jaccard dissimilarity including all eDNA samples. **B** NMDS of Bray-curtis dissimilarity including all eDNA samples. **C** NMDS of Jaccard dissimilarity with the three outlying samples are removed. **D** NMDS of Bray-Curtis dissimilarity with the three outlying samples are removed (Same data as Figure 5D).
